# The Arabidopsis N-terminal Acetyltransferase NAA50 Regulates Plant Growth and Defense

**DOI:** 10.1101/2020.01.02.893115

**Authors:** Matthew Neubauer, Roger W. Innes

## Abstract

Stress signaling in plants is carefully regulated to ensure proper development and reproductive fitness. Overactive defense signaling can result in dwarfism as well as developmental defects. In addition to requiring a significant amount of energy, plant stress responses place a burden upon the cellular machinery, which can result in the accumulation of misfolded proteins and endoplasmic reticulum (ER) stress. Negative regulators of stress signaling, such as *EDR1*, ensure that stress responses are properly suspended when they are not needed. Here, we describe the role of an uncharacterized N-terminal acetyltransferase, NAA50, in the regulation of plant development and stress responses. Our results demonstrate that NAA50, an interactor of EDR1, plays an important role in regulating the tradeoff between plant growth and defense. Plants lacking *NAA50* display severe developmental defects as well as induced stress responses. Reduction of *NAA50* expression results in arrested stem and root growth and senescence. Furthermore, our results demonstrate that *EDR1* and *NAA50* are required for suppression of ER stress signaling. This work establishes that *NAA50* is essential for plant development and the suppression of stress responses, likely through the regulation of ER stress. These experiments demonstrate a role for N-terminal acetylation in the suppression of ER stress, as well as the tradeoff between stress responses and development.

**One Sentence Summary:** Knockout in Arabidopsis of the broadly conserved N-terminal acetyl transferase NAA50 induces ER stress, leading to severe dwarfism and induction of defense responses.

## Introduction

As sessile organisms, plants frequently encounter and respond to stress conditions such as drought, salinity, heat, and microbial infection. Various adaptations enable plants to defend themselves against these stresses, however, they often come at a significant cost (Cipollini et al., 2014). Plant defense responses require significant sacrifices by infected cells and tissues, which can negatively impact plant growth. The Hypersensitive Response (HR), a form of programmed cell death, is a primary mode of defense for infected plant cells (Greenberg and Yao, 2004). Thus, plants must carefully tailor their defense responses to conserve energy for growth and reproduction (Huot et al. 2014). This tradeoff is exhibited by enhanced resistance mutants such as *snc1-1* and *cpr1* which have constitutively active defense responses and are dwarfed (Li et al. 2001; Bowling et al., 1994).

Stress responses place strain upon the cellular machinery, which can result in endoplasmic reticulum (ER) stress (Bao and Howell, 2017). ER stress can occur during biotic or abiotic stress, as well as during normal developmental processes that place increased demands on the protein translation and protein secretion machinery (Vitale and Boston, 2008). Response to ER stress is mediated by the unfolded protein response (UPR), which occurs in two phases. The first phase aims to alleviate ER stress through increased expression of chaperones, removal and degradation of misfolded proteins from the ER, and reduction of protein translation (Williams et al., 2014; Liu and Howell, 2010). If these attempts are unsuccessful, the UPR transitions into a pro-apoptotic phase (Woehlbier and Hetz, 2011; Walter and Ron, 2011; Srivastava et al., 2018). Recent studies have demonstrated that UPR genes are required for plant growth and development (Kim et al., 2018; Bao et al., 2019). On the other hand, mutations that constitutively activate the UPR cause dwarfism (Iwata et al., 2018). Just as stress responses to external stimuli must be regulated to ensure proper growth and development, so must responses to internal stress and the UPR.

We have previously identified and characterized the *EDR1* gene and demonstrated its role in negatively regulating plant stress response signaling (Frye and Innes, 1998; Christiansen et al., 2011; Serrano et al., 2014). In particular, *EDR1* negatively regulates the salicylic acid (SA) and ethylene pathways (Frye et al., 2001; Tang et al., 2005). Mutant *edr1* plants display enhanced sensitivity to a variety of stimuli, including drought, pathogen infection, abscisic acid (ABA), and ethylene (Frye and Innes, 1998; Frye et al., 2001; Tang et al., 2005; Wawrzynska et al., 2008). The variety of *edr1*-related phenotypes implies that *EDR1* function impacts a diversity of plant stress responses. Interestingly, *edr1* plants appear phenotypically wildtype in the absence of external stresses. This transitory requirement of *EDR1* indicates that it is functionally active only after a stress response has been induced.

There remain many unanswered questions regarding EDR1 function. EDR1 is believed to negatively regulate KEG, an E3 ubiquitin ligase required for post-embryonic development and endomembrane trafficking (Wawrzynska et al., 2008; Gu and Innes, 2011; Gu and Innes, 2012). However, it is unclear whether EDR1 itself is a regulator of development or endomembrane trafficking. Interestingly, EDR1 primarily localizes to the ER, yet no ER-associated function of EDR1 has been demonstrated (Christiansen et al., 2011).

To gain a greater understanding of EDR1 function, we performed a yeast two-hybrid screen to identify potential substrates of EDR1. These screens yielded a particularly interesting hit, At5g11340, a predicted N-terminal acetyltransferase (NAT) that bears similarity to the human Naa50 protein (Fig. 1A).

**Fig. 1.**
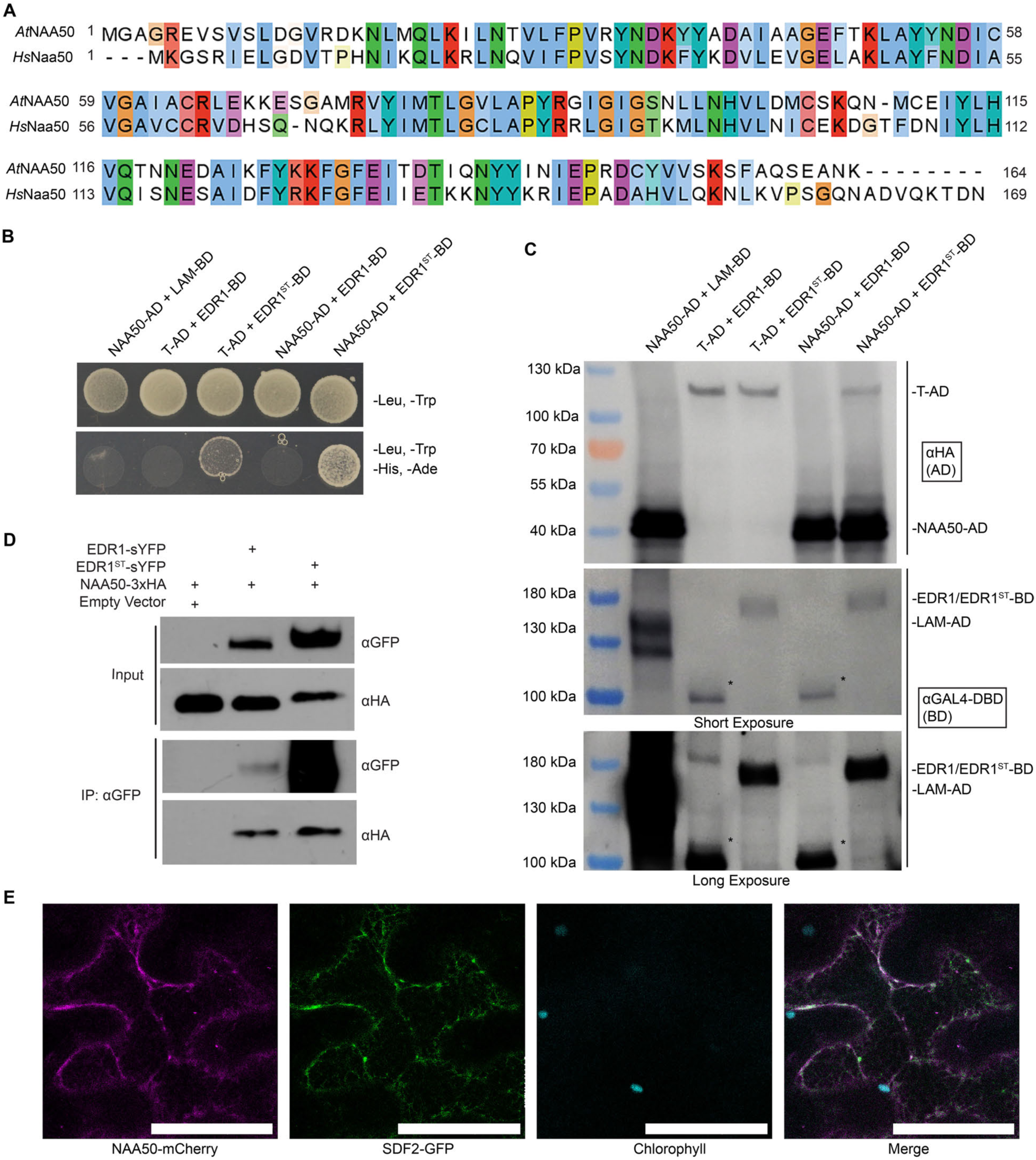
NAA50 physically interacts with EDR1. **A,** Naa50 is conserved in *Arabidopsis*. Amino acid alignment depicting *Arabidopsis* NAA50 and human Naa50. This alignment was generated using Clustal Omega (https://www.ebi.ac.uk/Tools/msa/clustalo/) and visualized in Jalview (Waterhouse et al., 2009). **B,** EDR1 interacts with NAA50 in yeast two-hybrid. AD, GAL4 activation domain fusion; BD, GAL4 DNA binding domain fusion. **C,** Immunoblot analysis of yeast strains from panel B. EDR1-BD accumulated poorly in yeast, and a significant accumulation of degraded EDR1-BD (*) was visible. **D,** NAA50 co-immunoprecipitates with EDR1. The indicated constructs were transiently expressed in *N. benthamiana* and then immunoprecipitated using GFP-Trap beads. **E,** NAA50 co-localizes with the ER marker SDF2. mCherry-tagged NAA50 and GFP-tagged SDF2 were transiently co-expressed in *N. benthamiana*. Bars = 50 microns. These experiments were repeated three times with similar results.

NATs serve as the catalytic components of larger complexes, designated as NatA-F in humans (Reviewed in Polevoda et al., 2009; Aksnes et al., 2016). Human Naa50 serves as the catalytic component of the NatE complex, which also includes the Naa10 and Naa15 subunits (Arnesen et al., 2006). Naa10, Naa15, and Naa50 are also found in the NatA complex, for which Naa10 provides catalytic function. NAT complexes mediate N-terminal acetylation (NTA), a widespread co-translational protein modification believed to affect the majority of eukaryotic proteins (Brown and Roberts, 1976; Polevoda and Sherman, 2003; Arnesen et al., 2009). These complexes target unique N-terminal sequences. Human Naa50 preferentially targets N-termini that have retained their initiator methionine and have a hydrophobic residue in the second position (Evjenth et al., 2009; Van Damme et al., 2011).

Based on work in yeast and humans, there is a solid biochemical understanding of how NATs function; however, the purpose of NTA is not well understood. Emerging evidence suggests that NTA serves various functions. In humans, the Golgi-localized Naa60 specifically targets transmembrane proteins and is required for the maintenance of Golgi integrity (Aksnes et al., 2015). Recent work in plants has implicated NTA in the regulation of stress responses and development. Both *NAA10* and *NAA15* are essential for plant embryonic development, and knockdown of either results in morphological defects and drought resistance (Feng et al., 2016; Linster et al., 2015). Differential NTA of the SNC1 receptor was found to have significant impacts on its activity, demonstrating a role for NTA in the regulation of defense signaling (Xu et al., 2015). Plant NATs bear strong similarity to their non-plant orthologues; however, the discovery of the plant-specific, plastid-localized NatG indicates that NTA in plants may serve unique purposes (Dinh et al., 2015). This early work demonstrates that NTA plays an important role in plant physiology and stress responses. However, many aspects of plant NATs have yet to be investigated.

Here, we demonstrate a role for the uncharacterized *Arabidopsis NAA50* gene in regulating plant growth and stress responses. Using knockout and transgenic knockdown lines, we show that *NAA50* is indispensable for normal plant growth and development. Loss of *NAA50* triggers defense response pathways in *Arabidopsis*, implicating *NAA50* in the negative regulation of defense signaling. Loss of *NAA50* also induces constitutive ER stress, while loss of *EDR1* leads to enhanced sensitivity to ER stress. Thus, both *EDR1* and *NAA50* appear to be involved in the negative regulation of ER stress. This work demonstrates the importance of NTA in plant stress responses and development, as well as a potential link between NTA and ER stress.

## Results

### NAA50 Interacts with EDR1

To verify the initial yeast two-hybrid screen which identified NAA50 as a potential interactor of EDR1, we performed additional assays to detect protein-protein interaction. To test for physical interactions between EDR1 and potential substrates, we utilized a “substrate-trap” mutant of EDR1, EDR1^ST^ (Gu and Innes, 2011). EDR1^ST^ contains a D810A substitution in the phosphotransfer domain, which is necessary for substrate phosphorylation, thus stabilizing the potential interaction between EDR1 and its substrates (Gibbs and Zoller, 1991). Our initial yeast two-hybrid screen was carried out using EDR1^ST^ as bait. In yeast two-hybrid, NAA50 was found to physically interact with EDR1^ST^, but not wildtype EDR1 (Fig. 1B). This result indicates that NAA50 may be a substrate of EDR1. However, immunoblotting demonstrated that wildtype EDR1 accumulation is significantly lower than that of EDR1^ST^ in yeast, potentially explaining the absence of an interaction (Fig. 1C). Co-immunoprecipitation using proteins expressed transiently in *N. benthamiana* demonstrated that NAA50 physically associates with both EDR1 and EDR1^ST^ *in vivo*, contrasting with our yeast two-hybrid results (Fig. 1D). EDR1 has been previously demonstrated to localize to the ER (Christiansen et al., 2011). We similarly observed an ER localization of NAA50 tagged with mCherry when transiently expressed in *N. benthamiana* (Fig. 1E). NAA50 co-localized with the GFP-tagged ER marker SDF2 (Nekrasov et al., 2009). These experiments indicate that EDR1 and NAA50 physically interact, that both proteins localize to the ER, and that NAA50 may be a substrate of EDR1.

### *Arabidopsis* NAA50 is Highly Conserved and Essential for Development

The discovery that NAA50 physically interacts with EDR1 prompted us to investigate its potential functions in *Arabidopsis*. There is a 51.25% identity match between *Arabidopsis* and human NAA50 proteins (Fig. 1A). This high degree of sequence similarity indicates that NAA50 function is likely conserved between plants and animals.

To investigate the role of NAA50 in plants, we characterized two T-DNA insertion mutants (SAIL_210_A02 and SAIL_1186_A03), which we designated *naa50-1* and *naa50-2*. Both mutant lines were found to be severely dwarfed compared to wild-type plants (Fig. 2, A–B). Knockout *naa50* seedlings displayed abnormal and dwarfed growth (Fig. 2A). As they developed, *naa50* plants remained dwarfed and were sterile, although stems and flowers did form (Fig. 2C). We were able to fully complement the *naa50-1* mutant phenotypes by transformation of a transgene carrying NAA50 tagged with sYFP under the control of the native NAA50 promoter, demonstrating that loss of NAA50 is responsible for the dwarf phenotype and sterility (Fig. 2B). These observations establish that *NAA50* is essential for normal plant growth and development.

**Fig. 2.**
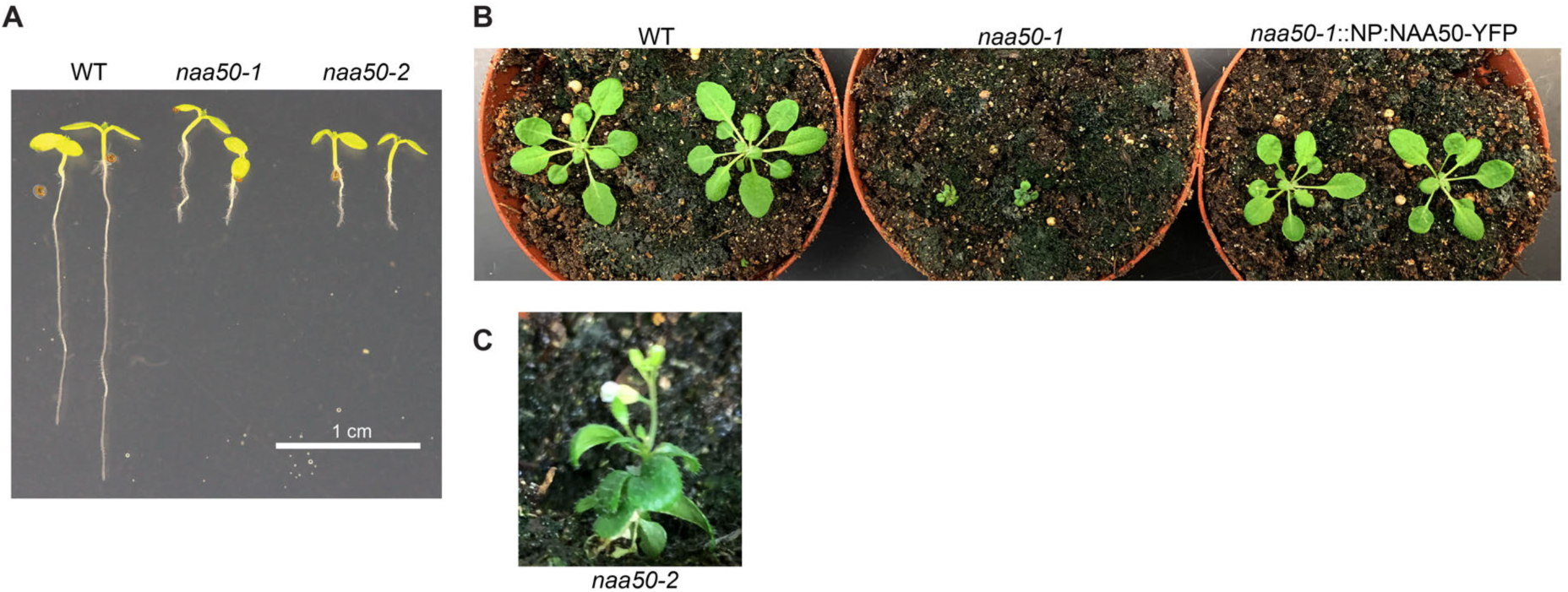
*NAA50* is required for plant development. **A,** Loss of *NAA50* results in dwarfed seedlings. Representative seven-day-old, MS-grown seedlings are depicted. **B,** *NAA50*-sYFP complements *naa50*-mediated dwarfism. Four-week-old adult plants are shown. NP, Native *NAA50* Promoter. **C,** *naa50* plants can develop stems and flowers. A five-week-old *naa50-2* plant is shown.

### Loss of Naa50 Alters Plant Growth

In addition to being dwarfed, *naa50* seedlings displayed a variety of developmental phenotypes. Root hair growth in *naa50* plants was irregular, and root hairs were elongated (Fig. 3A). This led us to hypothesize that loss of NAA50 may result in altered vacuole development. Loss of KEG, another EDR1-interacting protein, has been shown to result in altered vacuolar development (Gu and Innes, 2012). In *naa50-1* seedlings expressing the tonoplast marker γTIP (Nelson et al., 2007), altered vacuole shape was observed (Fig. 3B). Many *naa50-1* vacuoles appeared fractured and contained many “blebs”, similar to those observed in *keg* mutant seedlings (Gu and Innes, 2012). Additionally, *naa50-1* root cells were larger and irregularly shaped. This could indicate that NAA50 is involved in vacuole maturation.

**Fig. 3.**
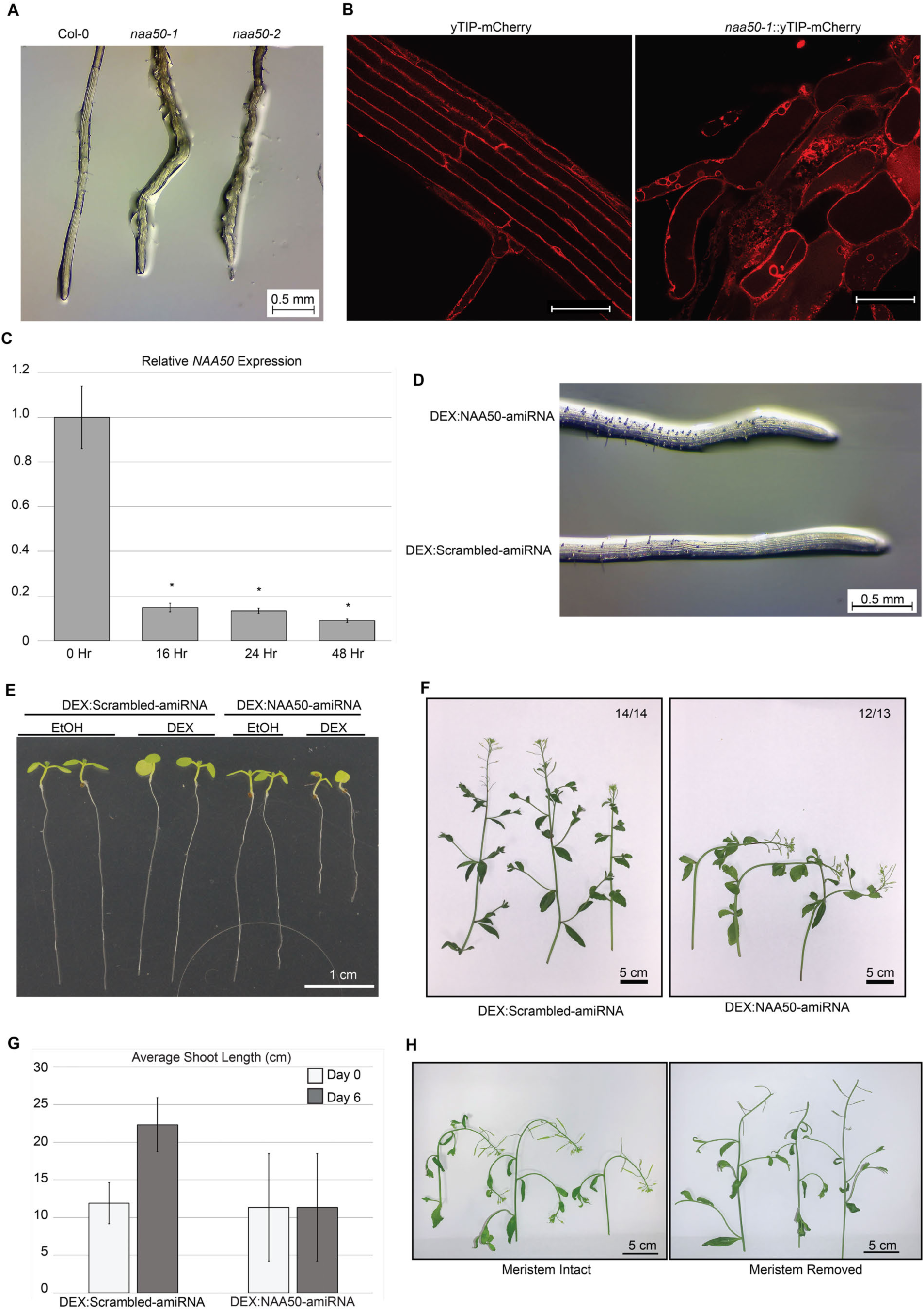
Loss of *NAA50* results in developmental changes. **A,** *naa50* seedlings have altered root morphology. The seedling roots depicted are from one-week-old seedlings. **B,** Vacuole and cell morphology are altered in *naa50* seedling roots. Shown are fluorescence micrographs taken of seven-day-old wildtype and *naa50-1* seedlings expressing mCherry-tagged γTIP. Scale bars = 50 microns. **C,** Dexamethasone treatment induces knockdown of *NAA50* in DEX:NAA50-ami plants. q-RT PCR was performed on cDNA generated from multiple adult DEX:NAA50-ami plants following dexamethasone treatment. Displayed are the averages of three replicates. Asterisk denotes *P* value < 0.05. Expression values were normalized to *ACTIN2*. This experiment was repeated three independent times with similar results. **D,** *NAA50* knockdown induces changes to root cell morphology. Five-day-old seedlings were transferred from MS plates to MS plates supplemented with DEX. Images were taken three days after dexamethasone exposure. **E,** *NAA50* knockdown slows root elongation. Seven-day-old seedlings were transferred to MS plates supplemented with ethanol or dexamethasone. Images were taken three days after transfer to ethanol- or dexamethasone-supplemented media. **F,** *NAA50* knockdown induces stem bending. Images were taken 24 hours after dexamethasone treatment. Numbers indicate proportion of all stems which displayed the given morphology. **G,** *NAA50* knockdown stalls stem growth. Stem measurements were taken on DEX:Scrambled-ami (n = 8) and DEX:NAA50-ami (n = 10) immediately before and six days after dexamethasone treatment. No stem growth was detected in DEX:NAA50-ami plants. **H,** Removal of the apical meristem inhibits *NAA50* knockdown-mediated stem bending. Adult DEX:NAA50-ami plants were sprayed with dexamethasone and images were taken twenty-four hours later. The shoot apical meristem was removed immediately prior to dexamethasone treatment.

The severe dwarfing and sterility of *naa50-1* homozygous mutant plants compromised our ability to study the role of *NAA50* in later stages of plant development. To overcome this limitation, we generated inducible knockdown plants based on the expression of an artificial microRNA (amiRNA) driven by a dexamethasone-inducible promoter (DEX:NAA50-ami). We identified two independent transgenic lines carrying this construct which displayed a significant knockdown of *NAA50* as early as 16 hours after dexamethasone treatment (Fig. 3C). As a control, we utilized a scrambled amiRNA line (DEX:Scrambled-ami), which contains a dexamethasone-inducible amiRNA with no predicted targets.

Knockdown of *NAA50* in the DEX:Naa50-ami plants resulted in severe morphological changes. Growth of DEX:NAA50-ami seedlings on MS media supplemented with dexamethasone increased the length of root hairs, recapitulating the *naa50* root hair phenotype (Fig. 3D). Additionally, dexamethasone treatment caused DEX:NAA50-ami seedlings to grow significantly slower than the control lines, resulting in shorter roots (Fig. 3E). *NAA50* knockdown also elicited changes in stem growth. 24 hours after dexamethasone treatment, the stems of DEX:NAA50-ami plants bent approximately 90° (Fig. 3F). As in the roots, dexamethasone treatment completely halted any growth of the primary stem in DEX:NAA50-ami plants (Fig. 3G). Interestingly, this shoot bending phenotype was suppressed by removal of the shoot apical meristem prior to dexamethasone treatment (Fig. 3H), suggesting that the bending phenotype is dependent on auxin redistribution. Our observations of knockout and knockdown plants confirm that *NAA50* is essential for normal plant growth and development.

### Loss of Naa50 Triggers Cell Death

As well as inducing growth changes, knockdown of *NAA50* caused early senescence in leaves. Leaves of adult DEX:NAA50-ami plants turned yellow and became necrotic following dexamethasone treatment (Figure 4A). Senescence also occurred in DEX:NAA50-ami seedlings after transfer to MS plates supplemented with dexamethasone (Fig. 4B). In both adults and seedlings, the senescence phenotype developed about 4 days after the initial dexamethasone treatment, long after the changes in growth rate and stem bending occurred.

**Fig. 4.**
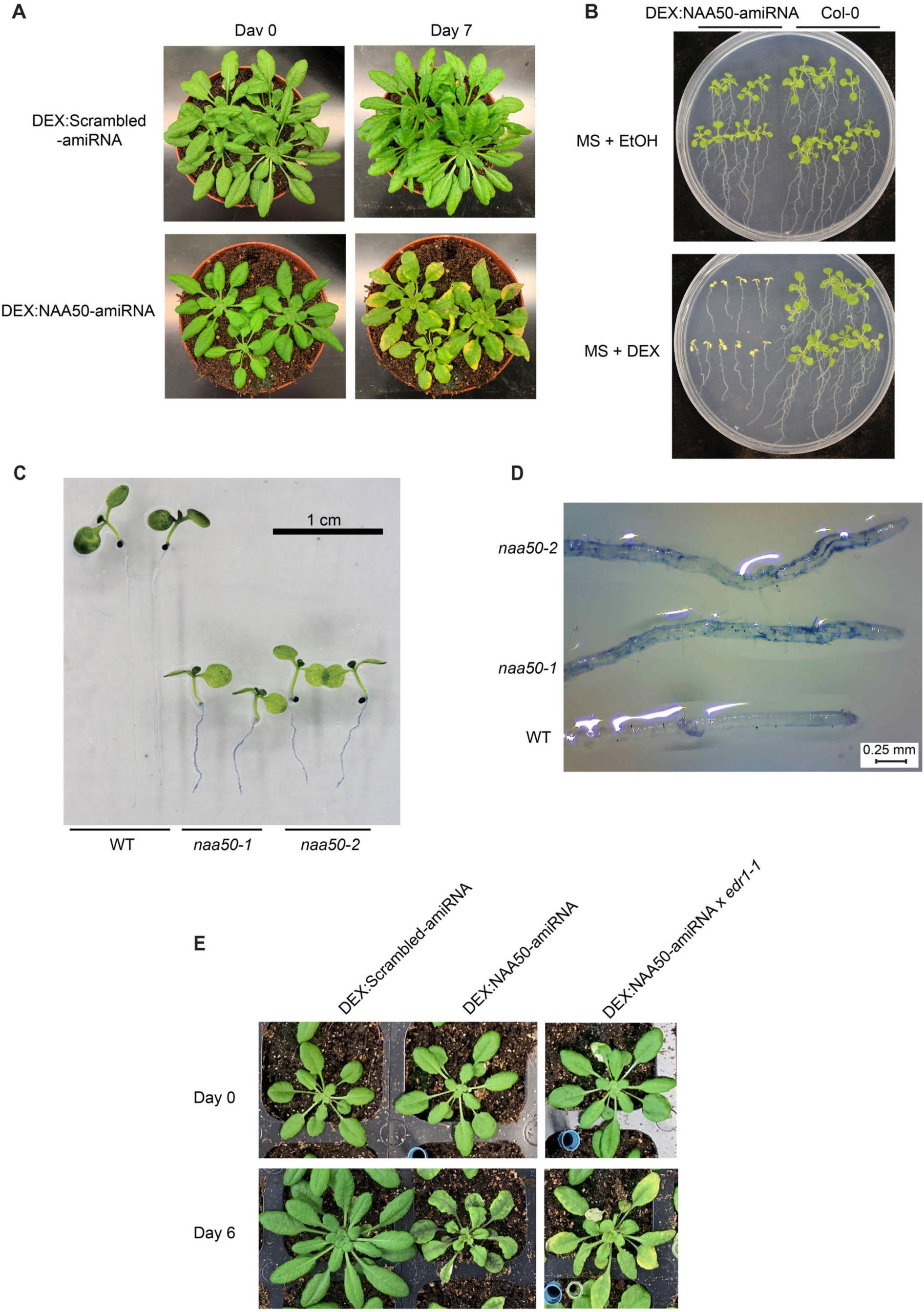
Loss of *NAA50* induces cell death and senescence. **A,** *NAA50* knockdown induces senescence in adult leaves. Four-week-old plants were sprayed with dexamethasone. Images were taken immediately before, and seven days after treatment. **B,** *NAA50* knockdown induces senescence in seedlings. Seedlings were grown on MS plates for seven days, and then transferred to MS plates supplemented with ethanol or dexamethasone. Images were taken seven days after transfer to ethanol- or dexamethasone-supplemented media. **C,** *naa50* seedling roots contain dead cells. Seven-day-old seedlings were stained with trypan blue dye. **D,** Cell death staining in *naa50* roots is spotty and irregular. Images depict trypan blue-stained roots from seven-day-old seedlings. **E,** Loss of *EDR1* does not alter senescence in *NAA50* knockdown plants. Images were taken of four-week-old plants immediately before, and seven days after dexamethasone treatment.

The discovery that knockdown of *NAA50* induces cell death prompted us to investigate whether loss of *NAA50* results in cell death in *naa50-1* seedlings. Indeed, roots of *naa50-1* and *naa50-2* seedlings were readily stained by trypan blue dye, indicating that loss of *NAA50* leads to the accumulation of dead cells in roots (Fig. 4C). Trypan blue staining of *naa50* roots was spotty and irregular, indicating that only a subset of *naa50* root cells died (Fig. 4D). Taken together, these results demonstrate that in addition to being essential for plant development, *NAA50* is also required for the repression of cell death and senescence.

Given the interaction between EDR1 and NAA50, we hypothesized that introduction of the *edr1-1* allele into *NAA50* knockdown plants may affect the senescence phenotype. However, we did not observe any major change in the senescence phenotype when *edr1-1* was introduced (Fig. 4E). That the *edr1-1* mutation did not enhance or suppress the senescence phenotype indicates that *NAA50* and *EDR1* may regulate senescence through a shared mechanism.

### Loss of Naa50 Represses Growth and Induces Stress Signaling

The discovery that knockdown of *NAA50* triggers changes in plant growth and senescence prompted us to investigate the transcriptional changes taking place in these plants. We therefore conducted an RNA sequencing-based analysis of the DEX:NAA50-ami transcriptome. Four-week-old plants were treated with dexamethasone, and RNA was collected 0, 12, and 24 hours later. The scrambled amiRNA line was utilized as a control. This design enabled a comparison of the DEX:NAA50-ami transcriptome at various time points, while excluding potential off-target effects of dexamethasone treatment or amiRNA overexpression.

Our RNA sequencing analysis indicated that *NAA50* knockdown resulted in altered expression of approximately 2,000 genes by 12 hours post-dexamethasone application (Supplemental Datasets). To determine the biological processes most impacted by loss of *NAA50*, we analyzed the biological gene ontology (GO) term enrichment in the 12 and 24 hour DEX:NAA50-ami datasets. This analysis demonstrated that *NAA50* knockdown leads to upregulation of genes involved in stress hormone signaling, as well as biotic and abiotic stress responses, while causing downregulation of a variety of plant growth and photosynthetic processes (Fig. 5A). In particular, transcripts of genes involved in photosynthesis, light responses, and growth hormone responses were negatively impacted. These changes in expression correlate with the altered development and induced senescence phenotypes observed during *NAA50* knockdown.

**Fig. 5.**
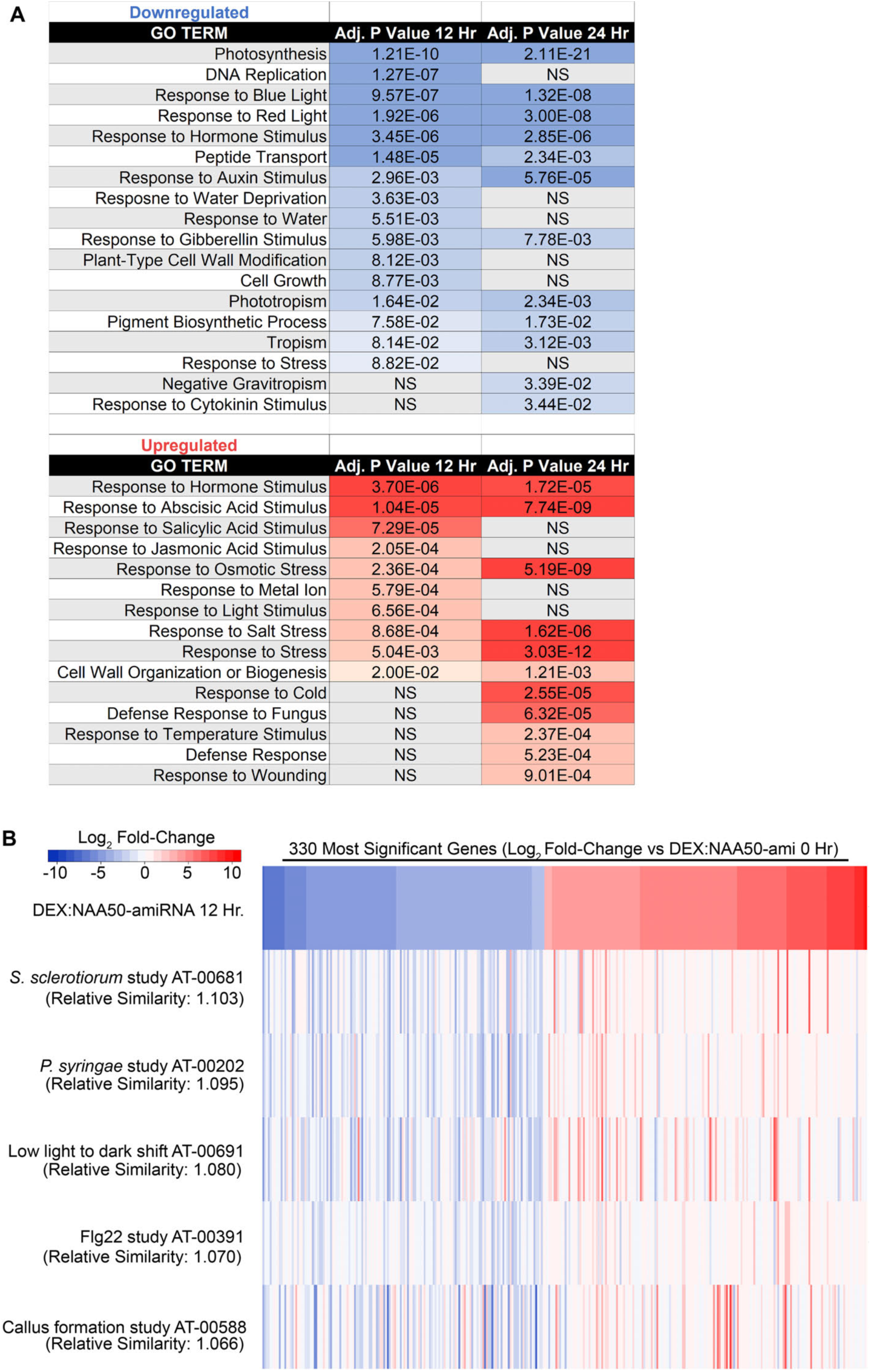
*NAA50* knockdown induces changes to growth and defense signaling. **A,** *NAA50* knockdown results in a downregulation of growth signaling, and an upregulation of defense signaling. Gene Ontology (GO) term enrichment analysis was performed using the BiNGO application to determine whether the DEX:NAA50-ami transcriptome was enriched for specific biological processes. NS, not statistically significant. **B,** The DEX:NAA50-ami transcriptome bears similarity to biotic and abiotic stress studies. The 330 most significantly altered transcripts (based on Log_2_ fold-change) from the DEX:NAA50-ami 12 hour dataset were compared to previous studies using the Genevestigator Signature tool. The five most related transcriptomes based on the calculated Relative Similarity scores are shown. A heatmap was generated using Heatmapper (http://www2.heatmapper.ca/expression/) to display the relative log_2_ fold-change for each of the 330 transcripts for each study.

To further analyze our transcriptome data, we searched for studies that had identified similar transcriptional changes using the Genevestigator Signature tool (Hruz et al., 2008). We selected the most significantly altered transcripts within the 12 hour DEX:NAA50-ami dataset, and searched for studies that displayed similar expression profiles. We found that the most similar expression profiles were those of studies investigating plant pathogen interactions, or light stress (Fig. 5B). This overlap demonstrates that *NAA50* knockdown elicits stress signaling in plants.

### *Naa50* and *EDR1* Repress ER Stress

We have previously found that plants lacking *EDR1* display an enhanced ER stress phenotype (unpublished). To verify this, we tested *edr1-1* plants for ER stress sensitivity by injecting leaves with tunicamycin (TM), an inhibitor of protein glycosylation that induces ER stress. Injected regions of *edr1-1* leaves senesced more rapidly than wild-type leaves (Figure 6A). This observation suggests that *EDR1* is required for proper execution of the unfolded protein response, or that loss of *EDR1* results in enhanced cell death signaling during ER stress signaling.

**Fig. 6.**
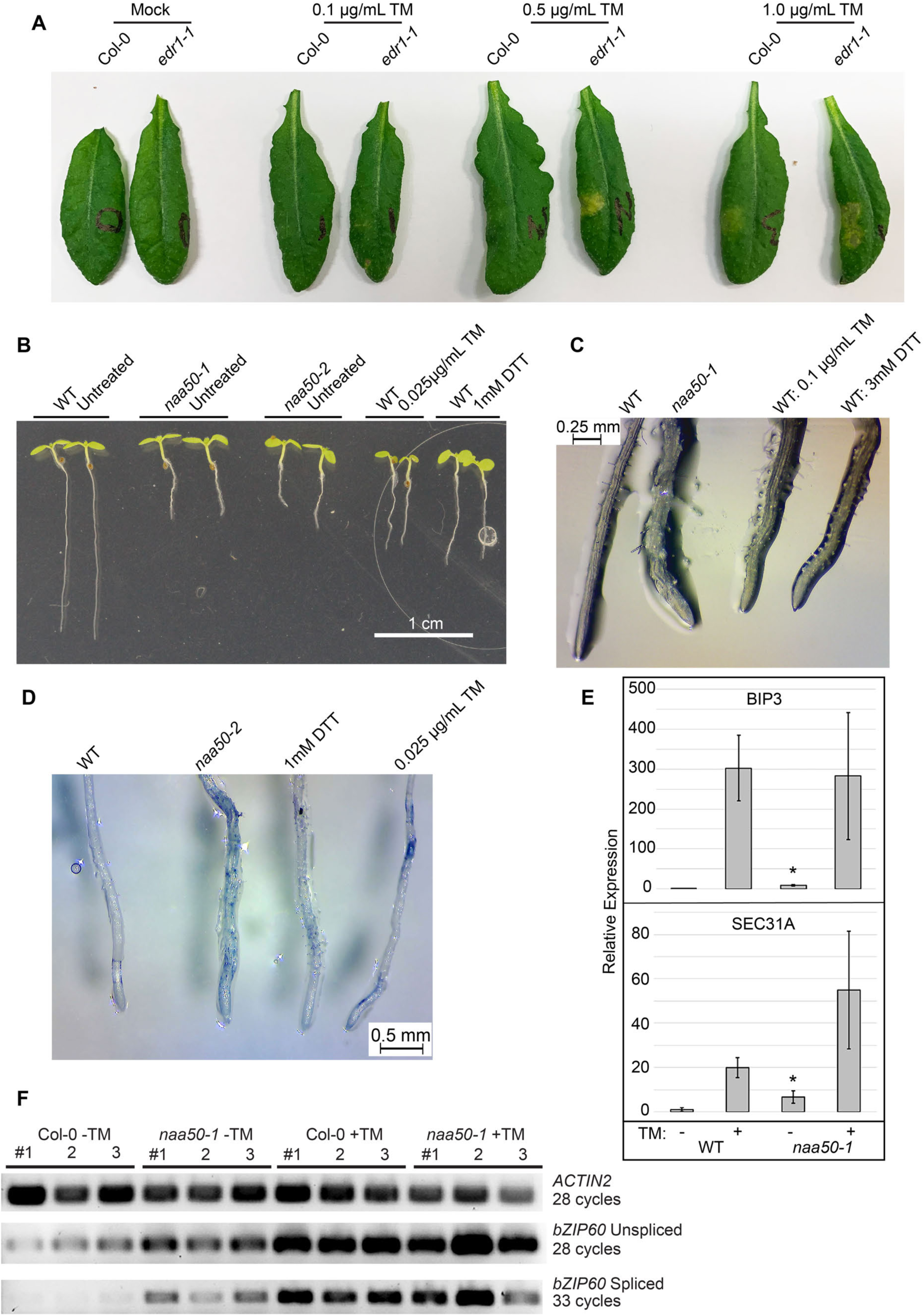
Loss of *EDR1* and *NAA50* result in changes to ER stress signaling. **A,** *edr1-1* mutants display heightened ER stress sensitivity. Leaves from six-week-old plants were infiltrated with various concentrations of tunicamycin using a needleless syringe. Leaves were removed, and images taken three days after injection. **B,** ER stress induces *naa50-*like root dwarfism. Seedlings were germinated on MS plates or MS supplemented with TM or DTT. Representative ten-day-old seedlings are shown. **C,** ER stress induces *naa50-*like root cell morphology. Roots of ten-day-old seedlings are depicted. Seedlings were grown on regular MS plates or MS plates supplemented with TM or DTT. **D,** ER stress induces cell death in roots. Ten-day-old seedlings were stained with trypan blue after growth on MS or MS supplemented with TM or DTT. **E,** *naa50-1* seedlings display heightened ER stress signaling in the absence of TM treatment. qRT-PCR was performed on cDNA generated from wildtype and *naa50-1* seedlings. Seedlings were germinated on MS plates and 5 days later transferred to regular MS or MS supplemented with 1 μg/mL TM. RNA was collected twenty hours after transfer to new plates. Gene expression values were normalized to *ACTIN2*. Values depict the averages of three biological replicates, each consisting of twenty individual seedlings. Error bars represent standard deviation between three independent biological replicates. Asterisk denotes *P* value < 0.05. **F,** *bZIP60* splicing is induced in *naa50-1* seedlings. RT-PCR was performed on the same cDNA used in panel E. Each lane represents a unique biological replicate derived from twenty seedlings.

NTA has been shown to alter protein stability, localization, and transport (Arnesen, 2011). This raised the question of whether loss of NAA50-mediated NTA may lead to induction of ER stress. Indeed, many of the observed *naa50*-mediated developmental phenotypes, such as stunted growth and cell death, can be caused by ER stress. Treatment with TM or dithiothreitol (DTT), which reduces disulfide bonds and induces ER stress, resulted in shorter roots, increased root hair length, and altered cell morphology in wild-type seedlings (Fig. 6, B–C). Additionally, TM and DTT treatments resulted in root cell death like that observed in *naa50* seedlings (Fig. 6D). These results demonstrate that ER stress treatment and loss of *NAA50* produce similar physiological changes.

To test whether *naa50* seedlings display constitutive ER stress responses, we measured transcription of ER stress marker genes by qPCR. *naa50-1* seedlings were found to have significantly higher levels of *BIP3* and *SEC31A* expression in the absence of any treatment (Fig. 6E). When treated with TM, however, *naa50-1* seedlings displayed WT levels of *BIP3* and *SEC31A* expression. During ER stress, the transcription factor *bZIP60* undergoes splicing, leading to its activation (Deng et al., 2011). Thus, detection of the spliced form of *bZIP60* indicates an active ER stress response. Untreated *naa50-1* seedlings were found to contain significantly higher levels of spliced *bZIP60* relative to WT (Fig. 6F). However, WT levels of *bZIP60* splicing occurred in TM-treated *naa50-1*. These results demonstrate that loss of *NAA50* leads to constitutive ER stress, but not an increase in maximum ER stress response signaling. Thus, *EDR1* and *NAA50* both appear to play important roles in regulating ER stress in plants.

### Naa50 Enzymatic Activity is Required for Development

Given the high sequence conservation between *Arabidopsis* and human NAA50 proteins (Fig. 1A), we hypothesized that the enzymatic activity of NAA50 would be conserved. In addition to functioning as an N-terminal acetyltransferase, human Naa50 has been shown to be capable of auto-acetylation (Evjenth et al., 2009). We therefore tested NAA50 for auto-acetylation activity using recombinant NAA50 protein. *In vitro* auto-acetylation assays using recombinant NAA50 protein demonstrated that *Arabidopsis* NAA50 is indeed capable of auto-acetylation (Fig. 7A).

**Fig. 7.**
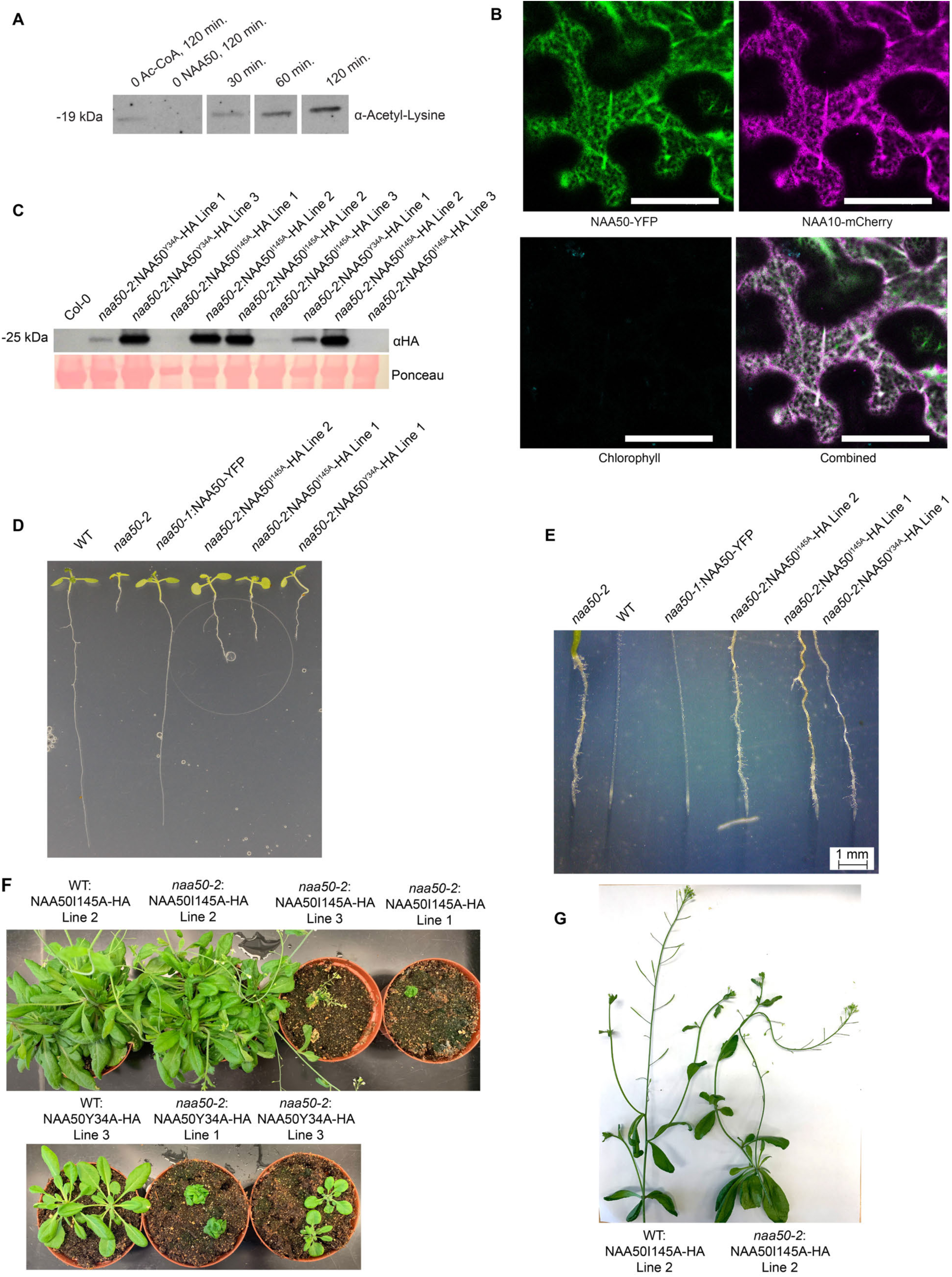
NAA50 enzymatic activity is required for plant development. **A,** Recombinant NAA50 displays auto-acetylation activity *in vitro*. Recombinant HIS-tagged NAA50 was expressed and purified from *E. coli. In vitro* reactions were performed at 30°C for the indicated time points. Samples were then boiled and subjected to gel electrophoresis and immunoblotting using an anti-acetyl-lysine antibody. This experiment was repeated three times with similar results. **B,** NAA50 co-localizes with NAA10. sYFP-tagged NAA50 was transiently co-expressed with mCherry-tagged NAA10 in *N. benthamiana.* Bars = 50 microns. **C,** Immunoblotting demonstrates that HA-tagged NAA50 mutant transgenes are expressed in transgenic plants. Leaf tissue from hygromycin-resistant T3 plants was subjected to gel electrophoresis and immunoblotting using an anti-HA antibody. **D,** Mutant *NAA50* transgenes do not complement *naa50* root dwarfism. Representative ten-day-old seedlings are depicted. **E,** Mutant *NAA50* transgenes do not complement *naa50* root cell morphology defects. Images were taken of representative ten-day-old seedlings. **F,** NAA50I145A can complement *naa50*-mediated rosette dwarfism. The top row depicts representative seven-week-old plants. The bottom row depicts representative 5-week-old plants. **G,** NAA50I145A does not complement *naa50-*mediated sterility. Stems were removed from 7-week-old plants.

Human Naa50 has previously been shown to associate with the NatA complex, which includes the Naa10 subunit (Arnesen et al., 2006). Transient expression of sYFP-tagged *At*NAA50 with mCherry-tagged *At*NAA10 indeed demonstrated that these proteins co-localize in plants (Fig. 7B).

Based on the sequence conservation between *Arabidopsis* NAA50 and human Naa50, as well as the co-localization of *At*NAA50 with *At*NAA10, we hypothesized that *At*NAA50 likely functions as an N-terminal acetyltransferase. To determine whether NAA50 is active in N-terminal acetylation, we tested whether various loss of function NAA50 mutants could complement *naa50-2* mutant phenotypes. *naa50-2* plants were transformed with NAA50^Y34A^-HA and NAA50^I145A^-HA. It has been demonstrated that the comparable Y31A and I142A mutations in human Naa50 reduce enzyme efficiency to below 10% and 42.2% of wild-type levels, respectively (Liszczak et al., 2011).

We were able to identify numerous transgenic lines expressing both the Y34A and I145A proteins (Fig. 7C). Following transformation with the NAA50^I145A^-HA transgene, we observed that the *naa50* root phenotype was not fully complemented in the transgenic lines, as roots retained their dwarf phenotype and altered cell morphology (Fig. 7, D-E). Despite retaining the *naa50* root phenotypes, some NAA50^I145A^ lines did not display the *naa50* dwarfism phenotype and had wildtype-sized rosettes (Fig. 7F). However, even when NAA50^I145A^ transgenic plants had wildtype-sized rosettes, they did not develop normal siliques or produce viable seed (Fig. 7G). The more severe NAA50^Y34A^ mutant also did not fully rescue *naa50-2* plants. NAA50^Y34A^ transgenic plants did not have normal roots or rosettes and were infertile (Fig. 7, D-F). Although the Y34A transgene was able to partially complement the rosette dwarfism, it was not able to fully complement the phenotype (Fig. 7F). For both the I145A and Y34A transgenic lines, we observed a correlation between NAA50 protein accumulation and rosette size (Fig. 7, C, F). The inability of NAA50^I145A^ and NAA50^Y34A^ transgenes to fully rescue *naa50-2* plants demonstrates the importance of NAA50-mediated NTA in plant growth and development. That the NAA50^I145A^ mutant was able to complement the rosette dwarfism, but not the root phenotypes or sterility demonstrates that NAA50-mediated NTA may be especially required for the growth and development of roots as well as fertility.

## Discussion

### *NAA50* is Required for Growth and the Suppression of Stress Responses

The investigation of NTA in regulating cell signaling in eukaryotes is still in its infancy, and identification and characterization of all plant NATs is incomplete. Our understanding of how NATs function comes primarily from work in human cell culture and yeast. However, recent work in plants has demonstrated a role for NTA in regulating diverse processes (Linster et al., 2015; Xu et al., 2015). Post-translational modification of proteins has long been appreciated as a mechanism by which cell signaling and crosstalk is regulated (Hunter, 2007). NTA may provide a mechanism by which plants regulate responses to external and internal stress signals at the translational level.

With this work, we have begun to characterize the role of *Arabidopsis NAA50* in regulating plant development and stress responses. Complete loss of *NAA50* results in severely dwarfed and sterile plants, as well as altered root morphology. By using hormone-inducible amiRNA transgenic plants, we demonstrated that *NAA50* knockdown results in reduced expression of developmental process and inhibits growth. Taken together, these results indicate that *NAA50* is required for plant growth and development.

Our work adds to growing evidence that NATs are required for plant development. *NAA10* and *NAA15* have previously been demonstrated to be essential for development. Loss of function mutations in *NAA10* or *NAA15* are embryonic lethal (Linster et al., 2015; Feng et al., 2016), while partial loss of *NAA15* results in dwarfism and enhanced defense signaling (Xu et al., 2015). Knockdown of *NAA10* and *NAA15* alters root morphology, enhances the growth of the primary root and inhibits the growth of lateral roots (Linster et al., 2015). Although NatA and NatE are required for proper development, loss of NatB is less severe (Ferrandez-Ayela et al., 2013; Xu et al., 2015). Similarly, loss of NAA30, the catalytic component of NatC, does not result in lethality in plants. It does, however, result in minor dwarfism as well as defects in photosystem II efficiency (Pesaresi et al., 2003). The range of developmental phenotypes resulting from mutations in NATs demonstrate that NATs differ in their involvement in plant development.

Our results demonstrate that loss of *NAA50* results in the activation of plant stress signaling. Knockdown of *NAA50* elicits senescence in adults as well as seedlings, while the roots of *naa50* seedlings contain an abundance of dead cells. Gene expression analysis confirmed that knockdown of *NAA50* results in an upregulation of defense signaling.

NATs appear to play unique roles in the regulation of plant stress responses. Loss of NatA has been shown to increase drought tolerance (Linster et al., 2015). The NatA and NatB complexes have been previously implicated in the regulation of the NLR protein SNC1 (Xu et al., 2015). Partial loss of *NAA15* results in increased stability and accumulation of SNC1, as well as enhanced defense signaling and resistance. Interestingly, loss of NatB leads to decreased accumulation of SNC1, and suppression of *snc1*-induced dwarfism (Xu et al., 2015). Our observations demonstrate that loss of NatE has a similar effect as the loss of NatA in plants, indicating that both are required for the suppression of defense signaling in the absence of external stress.

In addition to its role in negatively regulating defense signaling, our results implicate *NAA50* in the repression of ER stress. The developmental defects observed in *naa50* plants can be recapitulated by TM and DTT treatment, indicating that they may result from constitutive activation of ER stress responses. In support of this hypothesis, we observed increased expression of ER stress genes and *bZIP60* splicing in untreated *naa50* seedlings (Fig. 6). Following TM treatment, *naa50-1* seedlings displayed WT levels of ER stress signaling. Therefore, loss of *Naa50* induces ER stress signaling, but does not lead to greater induction during TM treatment. Additionally, the expression of *BIP3* and *SEC31A* in *naa50-1* seedlings was significantly lower in the absence of TM than during TM treatment. This indicates that the level of constitutive ER stress which occurs in *naa50-1* plants is significantly lower than that elicited by chemical treatment. Based on these results, we believe that NAA50 is required for the prevention of protein misfolding and aggregation, which contribute to ER stress. Plant NATs have not previously been demonstrated to play a role in the regulation of ER stress. Although NatE seems to be required for the repression of ER stress, it is possible that other NAT complexes may be required as well.

Our results demonstrate that NAA50-mediated NTA is likely required for plant development. Although we were unable to detect NAA50-mediated NTA *in vivo*, our complementation experiments demonstrate that the NAA50^I145A^ and NAA50^Y34A^ mutations, which inhibit NTA activity, prevent the NAA50 transgene from fully complementing *naa50* plants. This demonstrates an essential role for NAA50-mediated NTA in root development and fertility.

### NTA May Regulate ER Stress

The enzymatic function of human Naa50 has been demonstrated previously (Liszczak et al., 2011; Van Damme et al., 2011; Reddi et al., 2016; Evjenth et al., 2009). A high degree of conservation has been demonstrated for other NATs. For instance, human NatA can complement yeast NatA mutants (Arnesen et al., 2009). Based on the high level of sequence similarity between *Arabidopsis* and human Naa50, it is probable that enzymatic function is conserved. We were able to detect auto-acetylation of recombinant *Arabidopsis* NAA50 *in vitro*, demonstrating that it is indeed a functional acetyltransferase (Fig. 7). In addition, we found that mutations that alter NAA50 NTA activity prevent complementation of *naa50* mutant phenotypes (Fig. 7). As in other organisms, *Arabidopsis* NAA50 localizes primarily to the ER (Fig. 1, Fig. 7B). These similarities to other Naa50 proteins demonstrate that NAA50 likely functions as an NTA in plants.

There is evidence from human and yeast systems for the involvement of NATs in responding to ER stress and protein aggregates. NTA is known to contribute to protein stability, trafficking, and translocation to the ER (Arnesen, 2011; Forte et al., 2011). The NatA complex has been implicated in the regulation of protein aggregation (Arnesen et al., 2010). HYPK, a NatA component, has chaperone activity, and has been shown to inhibit the formation of protein aggregates (Raychaudhuri et al., 2007). Loss of NatA components in yeast results in compromised heat shock sensitivity and signaling, indicating a potential role for NTA in regulating heat shock (Gautschi et al., 2003; Das and Bhattacharyya, 2016). There is an established link between the UPR and heat stress responses in plants. Heat shock can induce protein aggregation and ER fragmentation (Richter et al., 2010). During heat stress, the UPR is activated and ensures proper reproductive development (Deng et al., 2011; Deng et al., 2016). We have demonstrated that loss of *NAA50* in plants results in constitutive ER stress, adding additional evidence that NTA is involved in the repression of protein aggregation and ER stress.

There is a well-established link between ER stress, the UPR, and defense signaling in plants. Plants carrying loss of function mutations in the stearoyl-ACP desaturase *SSI2* exhibit dwarfism, enhanced accumulation of ER stress marker BiP3, and higher PR-1 expression (Iwata et al., 2018; Kachroo et al., 2001). This mirrors the increased biotic and ER stress signaling observed in *naa50* plants. Mutants lacking UPR regulators IRE1 and bZIP60 display enhanced susceptibility to bacterial pathogens, demonstrating a link between the UPR and SA-based defense signaling (Moreno et al., 2012). If ER stress cannot be properly maintained, the UPR shifts into a cell death phase (Woehlbier and Hetz, 2011; Walter and Ron, 2011). A recent investigation of the transcriptional changes that occur during a prolonged UPR demonstrated that transcripts associated with biotic stress responses are elicited during the UPR (Srivastava et al., 2018). Biotic stress signaling is also impacted by the ER Quality Control (ERQC) pathway. The membrane-bound receptors upon which plant defense signaling relies undergo maturation through the ERQC pathway. Impairment of ERQC machinery can result in enhanced susceptibility, as the receptors required for pathogen recognition are unable to function (Tintor and Saijo, 2014). Thus, compromised ER integrity can hinder plant pathogen responses. Unsurprisingly, plant pathogens have been found to attack the host ER for their own benefit. The mutualistic fungus *Piriformospora indica* induces cell death using an ER stress-dependent mechanism, enabling its colonization of the *Arabidopsis* root (Qiang et al., 2012). The high degree of overlap between ER stress and biotic stress responses opens the possibility that the observed increase in stress signaling in *NAA50* knockout and knockdown plants results from changes to ER stress, rather than direct regulation of stress responses by *NAA50*.

A link between NTA and osmotic stress in plants has been recently proposed (Linster et al., 2015; Asknes et al., 2016). It was demonstrated that NatA knockdown plants display enhanced drought tolerance. Furthermore, levels of NatA-mediated NTA were shown to fluctuate in response to ABA treatment (Linster et al., 2015). Here, we have demonstrated that plant NATs may be required for proper protein folding and the repression of ER stress. There is a demonstrated link between ER stress and osmotic stress in plants. Overexpression of the chaperone BiP in tobacco and soybean results in enhanced drought tolerance (Valente et al., 2008). BiP expression in soybean was found to inhibit both ER- and osmotic stress-induced cell death (Reis et al., 2011). In wheat, treatment with Tauroursodeoxycholic Acid alleviates osmotic stress-induced cell death and ER stress signaling (Zhang et al., 2017). Strong osmotic stress alters root architecture and induces cell death through an ER stress-dependent mechanism (Duan et al., 2010).

If NTA is indeed required to prevent induction of ER stress, then the enhanced drought resistance of NAT-deficient plants may be an indirect result of changes to ER stress signaling. We have demonstrated that loss of *NAA50* alters root morphology, resulting in shorter roots, longer root hairs, and the accumulation of dead cells. Furthermore, these changes appear to be the result of constitutive ER stress. Constitutive ER stress resulting from reduction in NAT expression may lead to priming of stress responses, which ultimately results in a resistance phenotype. Thus, the enhanced drought tolerance of NatA knockdown plants observed by Linster et al. 2015 may result from enhanced ER stress and UPR signaling, rather than a direct effect on osmotic stress responses.

Although loss of *NAA50* has a significant impact on *Arabidopsis* development, it does not result in lethality, as in *Naa10 and Naa15* knockouts (Linster et al., 2015; Feng et al., 2016). This indicates potential redundancy for NAA50-mediated NTA, or that NAA50 is only essential for certain developmental processes. Human NatE, NatC, and NatF target a common set of N-terminal peptides (Aksnes et al., 2016). Given this overlap of function, other NATs may be capable of acetylating NatE targets in its absence. Although NatF has been characterized in humans, no ortholog of the NatF catalytic component Naa60 exists in yeast or *Arabidopsis*. A BLAST search using the human Naa60 sequence returns NAA50 as the most similar *Arabidopsis* protein. It is unclear whether *Arabidopsis* NAA50 can function similarly to NatF. NAA50 does not appear to have the same Golgi localization as human Naa60 (Aksnes et al., 2015), as we observed it primarily localizing to the ER (Fig. 1E). NatC may be able to fulfill some functions for NatE, however, loss of function mutations in *Arabidopsis* NatC are less severe than that of NatE, producing only minor dwarf phenotypes (Pesaresi et al., 2003).

Most work on NTA has been performed in unicellular organisms, making it impossible to study whether NATs display tissue-specific functions. There are likely to be differences in the expression patterns of NATs in different tissues. According to the BAR ePlant browser (http://bar.utoronto.ca/eplant), root expression of NAA50 and NAA10 is predicted to be the highest in the meristematic region. It is likely that some NAT complexes are specifically active at certain developmental periods, or in specific tissues. If other plant NAT complexes are indeed able to fulfill some functions of NAA50, it is also possible that they are only able to do so in certain tissues or at certain points in development, based upon their own expression profiles. The study of plant NATs has the potential to expose tissue- and development-specific NAT activity.

Loss of *NAA50* especially affected certain cell types and tissues. In *NAA50* knockdown plants, loss of *NAA50* led to reduced growth of both roots and stems. Furthermore, stem bending and altered root morphology were observed. The use of an inducible knockdown line enabled us to compare the effects of *NAA50* knockdown in new and old cells. Interestingly, phenotypes resulting from *NAA50* knockdown were mainly exhibited in newly developed cells. The sterility of *naa50* plants demonstrates that *NAA50* is required for reproductive as well as vegetative development. These observations demonstrate that NAA50 activity may be especially required by developing cells, or cells undergoing rapid growth and division.

If *NAA50* is indeed required for the regulation of ER stress, it follows that roots, shoots, and anthers would be especially impacted by its loss. There is evidence that plant vegetative and reproductive development require an intact UPR to manage ER stress. Roots have been shown to be particularly sensitive to ER stress (Cho and Kanehara, 2017). Significant changes in root and shoot development result from mutations in UPR genes, indicating that a functional UPR is essential for vegetative development (Deng et al., 2013; Kim et al., 2018; Bao et al., 2019). Mutations in UPR genes also have a significant impact on plant reproductive development (Deng et al., 2013; Deng et al., 2016). In fact, the UPR is constitutively active in anthers (Iwata et al., 2008). The requirement for UPR signaling in unstressed plants implies that ER stress occurs during normal development and must be managed by the UPR, or that UPR genes are involved in the direct regulation of developmental genes (Kim et al., 2018). A requirement for the UPR in development has long been demonstrated in animals. The rapid production of immunoglobulins by B cells is preceded by an upregulation of the UPR, which manages potential ER stress (van Anken et al., 2003). Roots, shoots, and anthers may rely upon UPR signaling due to the high level of protein translation which occurs in these tissues during development. That these tissues were indeed particularly affected by the loss of NAA50 demonstrates that NAA50 may be required for the management of ER stress which occurs during development.

### Model for EDR1 and NAA50 Regulation of ER Stress

Our initial interest in NAA50 was based on its physical interaction with EDR1. Indeed, the enhanced defense signaling observed in *NAA50* knockout and knockdown plants correlates with many *edr1* phenotypes. *EDR1* and *NAA50* also appear to play a role in the regulation of ER stress. *edr1* plants were found to have enhanced sensitivity to TM treatment, while loss of *NAA50* induced constitutive ER stress.

Our work indicates that EDR1 and NAA50 may be involved in the repression of ER stress (Fig. 8). Since NAA50 likely functions primarily in the NTA of target peptides, loss of NAA50 may result in the translation of proteins which lack a required N-terminal acetylation mark. Loss of NAA50-mediated NTA likely results in the misfolding, improper trafficking, or aggregation of proteins, ultimately producing ER stress. We have found that loss of EDR1 results in increased ER stress sensitivity. It is possible that EDR1 activates NAA50, perhaps during a stress event. Thus, when plants lacking EDR1 encounter stress, NAA50 would lack proper activation. The lack of NAA50-mediated NTA would therefore lead to mild ER stress, ultimately resulting in enhanced senescence and cell death. This model provides a potential explanation for the wide range of stimuli to which *edr1* plants display enhanced senescence and sensitivity.

**Fig. 8.**
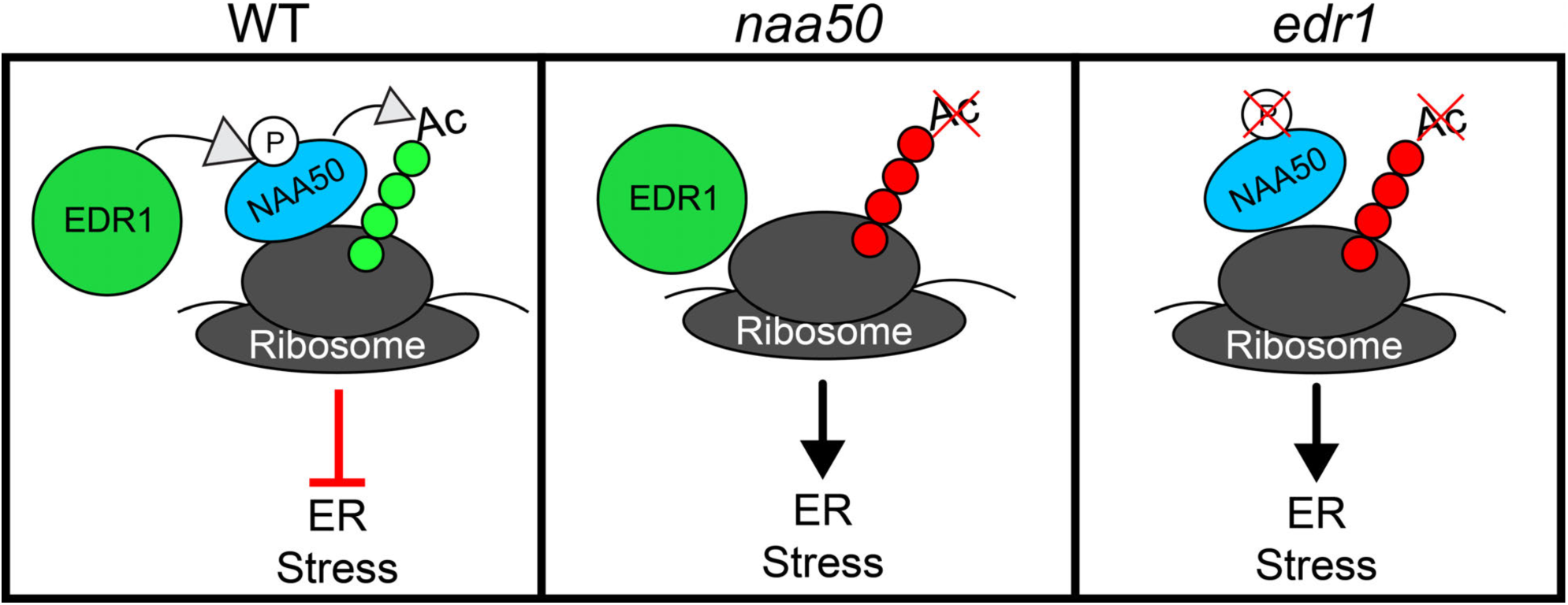
Model for EDR1- and NAA50-mediated regulation of ER stress. Left: In wildtype plants, EDR1 activates NAA50-mediated NTA, possibly through phosphorylation. NAA50-mediated NTA ensures proper protein folding, thereby inhibiting ER stress. Middle: In plants lacking functional NAA50, the absence of NAA50-mediated NTA results in protein aggregation and ER stress. Right: Biotic and abiotic stress events strain the translational machinery requiring altered or enhanced NAA50-mediated NTA. In plants lacking EDR1, NAA50 is not properly activated during these events, resulting in enhanced ER stress and senescence.

## Material and Methods

### Plant material and growth conditions

*Arabidopsis thaliana* accession Col-0, and Col-0 mutants *edr1-1* (Frye and Innes 1998), *naa50-1* (SAIL_210_A02), and *naa50-2* (SAIL_1186_A03) were used in this study.

For growth on Murashige and Skoog (MS) plates, seeds were surface sterilized with a solution of hydrogen peroxide and ethanol (1:19) and planted on one-half-strength MS plates supplemented with 0.8% agar. For soil-grown plants, seed was directly sowed onto Pro-Mix PGX Biofungicide plug and germination mix supplemented with Osmocote 14-14-14 fertilizer (ICL Fertilizers). Plates and flats were placed at 4°C for 48 hours for stratification before being transferred to a growth room set to 23°C and 12 hour light (150 µEm-2s-1)/12 hour dark cycle. For transient expression experiments, *Nicotiana benthamian*a was grown under the same growth room conditions as *A. thaliana*.

### Plasmid construction and generation of transgenic *Arabidopsis* plants

*NAA50* clones were derived by PCR amplification using cDNA from Col-0. Site-directed mutagenesis was utilized to introduce the I145A and Y34A mutations into *NAA50* (Qi and Scholthof, 2008). All primers used in this study for cloning and site-directed mutagenesis are listed in Supplementary Table S1.

For yeast-two hybrid assays, the full-length open reading frames of EDR1, EDR1 (D810A), and Lamin (LAM) were cloned into the DNA-binding domain vector pGBKT7 (Clontech Matchmaker System). The full-length open reading frame of NAA50, and the SV40 Large T Antigen (T) were cloned into pGADT7. EDR1 full-length wild-type cDNA and EDR1^ST^ (D810A) were cloned into pGBKT7 using SmaI and SalI restriction sites. NAA50 was cloned into pGADT7 using ClaI and XhoI restriction sites.

For transient expression in *N. benthamiana*, NAA50 was cloned into the cauliflower mosaic virus 35S promoter vector pEarleyGate100 (Earley et al. 2006) using a modified multisite Gateway recombination cloning system (Invitrogen) as described in (Qi et al. 2012). EDR1-sYFP and EDR1^ST^-sYFP were cloned into the dexamethasone-inducible pBAV154 (Vinatzer et al., 2006) using multisite Gateway cloning.

For the generation of amiRNA transgenic plants, a *NAA50*-specific amiRNA construct was created by PCR amplification following the procedures of Schwab et al. (2006), which included insertion of the *NAA50* sequence flanked by regions of the MIR319 microRNA. The resulting amiRNA construct was cloned into pBAV154 using Gateway cloning.

To generate transgenic plants containing NAA50-sYFP under the control of a native promoter, the 297 nucleotides upstream of the NAA50 start site were cloned into PMDC32 (Qi and Katagiri, 2009) using KpnI and HindIII restriction sites. NAA50^I145A^ and NAA50^Y34A^ were generated using site-directed mutagenesis (Qi and Scholthof, 2008), and cloned into pEarlyGate100 (Earley et al., 2006) with a C-terminal 3xHA tag using multisite Gateway cloning. Transgenic plants were generated using the floral dip method (Clough and Bent, 1998).

Plasmids were transformed into Agrobacterium strain GV3101 (pMP90) by electroporation with selection on Luria-Bertani plates containing 50 μg/mL kanamycyin sulfate (Sigma-Aldrich) and 20 μg/mL gentamicin (Gibco). Selection of transgenic plants carrying a BASTA resistance cassette was performed by spraying 1-week old seedlings with 300 μM BASTA (Finale), or by selection on MS plates supplemented with 300 μM BASTA. Selection of plants carrying a hygromycin resistance cassette was performed by germinating seed on MS plates supplemented with 20 µg/mL hygromycin (Fischer Scientific).

For expression in *E. coli*, NAA50 was cloned into pDEST17 using Gateway cloning. The resulting plasmid was transformed into *E. coli* strain BL21.AI (Invitrogen).

### Yeast two-hybrid assays

For yeast two-hybrid assays between EDR1 and NAA50, pGBKT7 and pGADT7 clones were transformed into haploid yeast strain AH109 (Clontech) by electroporation and selected on SD-Trp-Leu medium. Successful transformants were selected after 48 hours of growth at 30°C and then struck onto fresh SD-Trp-Leu medium and allowed to grow for another 48 hours. Before carrying out yeast two-hybrid assays, yeast was grown in liquid SD-Trp-Leu medium for 16 hours at 30°C. Cultures were re-suspended in water to an OD_600_ of 1.0, serially diluted, and plated on appropriate SD media. Plates were grown for up to 4 days at 30°C.

### Immunoprecipitations and immunoblots

For total protein extraction, tissue was ground in lysis buffer (50 mM Tris-HCl, pH 7.5, 150 mM NaCl, 0.1% Nonidet P-40, 1% Plant Proteinase Inhibitor Cocktail [Sigma], and 50 mM 2,2′-Dithiodipyridine [Sigma]) or, for co-IPs, IP Buffer (50 mM Tris, pH 7.5, 150 mM NaCl, 1mM EDTA, 0.1% Nonidet P-40, 10% glycerol, 1% Plant Proteinase Inhibitor Cocktail [Sigma], and 50 mM 2,2′-Dithiodipyridine [Sigma]). For expression of dexamethasone-inducible proteins, plants were sprayed with a 50 μM dexamethasone solution containing 0.02% (v/v) Silwet L-77 (OSi Specialties) 16 hours before tissue was harvested. Samples were centrifuged at 10,000 g at 4°C for 5 minutes, and supernatants were transferred to new tubes.

Immunoprecipitations were performed as described previously (Shao et al. 2003) using GFP-Trap_A (Chromotek). Total proteins were mixed with 1 volume of 2x Laemmli sample buffer, supplemented with 5% β-mercaptoethanol, 1% Protease Inhibitor Cocktail (Sigma), and 50 mM 2,2′-Dithiodipyridine (Sigma). Samples were then boiled for 5-10 minutes before loading. Total proteins and/or immunocomplexes were separated by electrophoresis on a 4-20% Mini-PROTEAN TGX Stain-Free protein gel (Bio-Rad). Proteins were transferred to a nitrocellulose membrane and probed with anti-HA-HRP (3F10) (Sigma), mouse anti-GFP (ab6556) (Abcam), and goat anti-mouse-HRP antibodies (A-10668) (Invitrogen).

For protein extraction from yeast, yeast grown on solid SD -Leu, -Trp plates were resuspended in lysis buffer (100 mM NaCl, 50 mM Tris-Cl, pH 7.5, 50 mM NaF, 50 mM Na-β-glycerophosphate, pH 7.4, 2 mM EGTA, 2 mM EDTA, 0.1% Triton X-100, 1 mM Na_3_VO_4_). Glass beads were then added to the suspension and the solution was vortexed for 1 minute three times. After the addition of 1 volume of 2x Laemmli sample buffer supplemented with 5% β-mercaptoethanol, samples were boiled for 10 minutes. Immunoblots were performed using anti-HA-HRP (3F10) (Sigma), mouse anti-GAL4DBD (RK5C1) (Santa Cruz Biotechnology), goat anti-mouse-HRP (A-10668) (Invitrogen), antibodies. Visualization of immunoblots from yeast strains used in two-hybrid assay were performed using the KwikQuant Imager (Kindle Biosciences).

### Fluorescence and light microscopy

Confocal laser scanning microscopy was performed on a Leica TCS SP8 confocal microscope (Leica Microsystems) equipped with a 63X, 1.2-numerical aperture water objective lens and a White Light Laser. sYFP fusions were excited at 514-nm and detected using a 522 to 545 nm band-pass emission filter. mCherry fusions were excited at 561 nm and detected using a custom 595 to 620 nm band-pass emission filter.

To capture detailed images of *Arabidopsis* roots, images were captured using a Stemi 305 compact Greenough stereo microscope (Zeiss). Digital images were captured using Labscope software (Zeiss).

### Quantitative-PCR

For RT-PCR and quantitative RT-PCR experiments, RNA was extracted using the Spectrum plant total RNA kit (Sigma-Aldrich) according to manufacturer’s instructions. cDNA was produced from 1 μg total RNA using the Verso cDNA synthesis kit (Thermo Fisher Scientific). Relative RNA amounts were determined by quantitative RT-PCR using the Power Up SYBR Green Master Mix (Thermo Fisher Scientific). A comparative Ct method was used to determine relative quantities (Schmittgen and Livak, 2008). ACTIN2 was used for normalization.

### *NAA50* knockdown transcriptome profiling

For RNA sequencing, plants were first sprayed with a solution containing 50 μM dexamethasone and 0.02% (v/v) Silwet L-77 (OSi Specialties) 24 hours, 12 hours, and immediately before tissue collection. Three biological replicates were performed per genotype per treatment, each consisting of approximately 0.4 g leaf tissue taken from the 4th leaf of 4 unique plants. RNA was extracted from 4-week-old Arabidopsis leaves using the Spectrum plant total RNA kit (Sigma-Aldritch) according to manufacturer’s instructions.

Total RNA was prepared into equimolar pools for each sample submitted to Indiana University’s Center for Genomics and Bioinformatics for cDNA library construction using a TruSeq Stranded mRNA LT Sample Prep Kit (Illumina) following the standard manufacturing protocol. Sequencing was performed using an Illumina NextSeq500 platform with 75 cycle sequencing kit generating 84bp single-end reads. After the sequencing run, demultiplexing was performed with bcl2fastq v2.20.0.422.

Trimmomatic (1; version 0.33; non-default parameters = ILLUMINACLIP:2:20:6 LEADING:3 TRAILING:3 SLIDINGWINDOW:4:20 MINLEN:35) was used to trim reads of adapter and low-quality bases. Reads were mapped to the *Arabidopsis thaliana* genome using STAR with the final parameters (4; version 2.5.2a; -- outSAMattributes All --outSAMunmapped Within --outReadsUnmapped Fastx -- outFilterMultimapNmax 1 --seedSearchStartLmax 25 --chimSegmentMin 20 --quantMode GeneCounts --twopassMode Basic --outWigType wiggle --outWigStrand Unstranded -- outWigNorm None --sjdbGTFtagExonParentTranscript Parent --sjdbGTFtagExonParentGene ID --outSAMtype BAM SortedByCoordinate). Read counts were determined using a custom perl script. Differential expression comparisons of all features with 5 or more reads (in total across all samples) were carried out with DESeq2 (2; R package version 3.4.0) along with the IHW (3) package to adjust for multiple testing procedures.

Identification of significantly altered transcripts was performed by comparing ‘DEX:NAA50-ami’ and ‘DEX:Scrambled-ami’ datasets. Transcripts which differed significantly (adjusted *P*-value < 0.05) between the ‘DEX:NAA50-ami’ and ‘DEX:Scrambled-ami’ datasets were then analyzed to determine whether expression had increased or decreased relative to the ‘DEX:NAA50-ami 0 hr’ dataset. Those transcripts which were significantly (adjusted *P-*value < 0.05 and log_2_ fold-change > 1.5) up- or downregulated relative to the ‘DEX:NAA50-ami 0 Hr’ dataset were then used for GO term enrichment analysis. Gene Ontology (GO) term enrichment analysis was performed in Cytoscape using the BiNGO app (Maere et al., 2005).

Transcriptome similarity analysis was performed using the Genevestigator Signature tool (https://genevestigator.com/gv/doc/signature.jsp). For this analysis, a list of the 330 most significantly altered (greatest log_2_ fold-change) transcripts at 12 hours was used as input. A heatmap comparing this input to similar transcriptomes was generated using the Heatmapper Expression tool (http://www2.heatmapper.ca/expression) (Babicki et al., 2016).

### Trypan blue staining

Trypan blue staining of *Arabidopsis* roots was performed by soaking seedlings in a solution of 10 mg/mL trypan blue (Sigma) in water for twenty minutes. Seedlings were then washed three times with deionized water.

### ER stress treatments

ER stress treatments of *Arabidopsis* seedlings were performed by growing seeds directly on MS plates supplemented with TM (Sigma) or DTT (Bio-Rad). For treatment of adult plants, TM was injected directly into one half of an *Arabidopsis* leaf using a needleless syringe.

### *In vitro* acetylation assays

*E. coli* strain BL21.AI was transformed with a pDEST17 vector carrying NAA50. 5xHIS-tagged NAA50 was purified from *E. coli* using a Nickel-His column (Sigma). A 5 mL culture was incubated at 37°C for 16 hours, and then subcultured to a final volume of 100 mL. The culture was grown until the OD_600_ reached 0.5. Expression of NAA50 was induced by adding 1 mM IPTG and 0.2% Arabinose to the culture. The culture was then incubated at 30°C for 3 hours. Cells were then harvested and resuspended in 8 mL of Native Purification Buffer (50 mM NaH2PO4, 500 mM NaCl) supplemented with 8 mg lysozyme and a protease inhibitor tablet (Roche). The suspension was then incubated on ice for thirty minutes, and then sonicated. After sonication, the cell debris was pelleted by centrifugation (5,000 x g, 15 minutes) at 4°C. The Ni-NTA resin was washed twice with Native Purification Buffer and then incubated with the lysate for 1 hour at 4°C. The resin was then washed 4 times with Wash Buffer (Native Purification Buffer supplemented with 6 mM Imidazole). Fractions were eluted with Elution Buffer (Native Purification Buffer supplemented with 250 mM Imidazole).

For *in vitro* auto-acetylation assays, 4 µg recombinant NAA50 was incubated with 100 µM Acetyl-Coenzyme A (Roche) in a 2X acetylation buffer (50mM Tris HCl pH 7.5, 2mM EDTA, 200mM NaCl, 10% Glycerol) at 30°C. Detection of auto-acetylation activity was performed through immunoblotting using acetylated lysine monoclonal antibody (1C6) (Invitrogen).

### Accession numbers

Arabidopsis sequence data is available under the following AGI accession numbers: EDR1 (At1g08720), NAA50 (At5g11340), γ-TIP (At2g36830), NAA10 (AT5G13780), SDF2 (AT2G25110).

## ACKNOWLEDGMENTS

We thank the Indiana University Light Microscopy Imaging Center for access to the Leica SP8 confocal microscope. We also thank James Ford and Doug Rusch from the Indiana University Center for Genomics and Bioinformatics for their work on the RNA sequencing experiment, and Reid Gohmann for assistance with ER stress assays on the *edr1* mutant.

